# Rapid and accurate quantification of viable *Listeria monocytogenes* with clonal specificity using microfluidic droplet digital PCR based technology

**DOI:** 10.1101/2025.11.27.690673

**Authors:** Chi Song, Zehang Gao, Gaoze Cai, Yongjie Yu, Ruihua Ding, Shilun Feng, Yangtai Liu

## Abstract

To overcome the technical bottlenecks in the precise quantification and molecular typing of viable foodborne pathogens, this study establishes a microfluidic droplet digital PCR (ddPCR) based method for rapid and accurate detection and quantification of viable *Listeria monocytogenes* with clonal specificity. In contrast to the time-consuming plate culture methods and unspecific rapid detection methods, the method in this study employs clonal complex (CC) specific primers and probe for strain-specificity and integrate the nucleic acid dye propidium monoazide (PMA) to effectively distinguish viable from dead bacteria. The rapid and precise quantification of viable bacteria is achieved through microdroplet counting. This method does not require DNA extraction, and the entire detection process takes only about 3 hours, with a quantitative detection limit of 3.3×10^2^ CFU/mL, providing strong technical support for the risk monitoring of highly pathogenic specific types of *L. monocytogenes*.

## 1. Introduction

Food safety is a critical issue concerning global public health and socio-economic development. One of the primary food safety hazards is foodborne disease caused by foodborne microorganisms [1]. These diseases severely threaten public health and hinder global economic development [2–5]. The World Health Organization estimates that approximately 600 million cases of foodborne illnesses occur worldwide each year, resulting in 420,000 deaths. Additionally, low- and middle-income countries suffer annual losses of $110 billion in productivity and medical expenses due to unsafe food [6]. *Listeria monocytogenes* is a facultative anaerobic foodborne pathogen that is widely present in the environment and food. It can survive under low temperatures, acidic environments, high salt concentrations, and other harsh conditions, often contaminating dairy products, vegetables, fish, meat products, and ready-to-eat foods [7]. Over 99% of human listeriosis cases are foodborne. High-risk groups susceptible to listeriosis include pregnant women, newborns, cancer patients, the elderly, and immunocompromised individuals, with a fatality rate as high as 20 ∼ 30% [8,9]. The European Food Safety Authority (EFSA) reported 2,952 confirmed cases of listeriosis caused by *L. monocytogenes* in Europe in 2023 [10]. According to the U.S. Centers for Disease Control and Prevention, *L. monocytogenes* causes approximately 1,600 cases of foodborne illness and 260 deaths annually [11]. In China, a total of 562 sporadic cases were reported from 2011 to 2017 [12]. Therefore, *L. monocytogenes* pose a threat to global food safety and human health [13].

Currently, the internationally recognized gold standard for detecting viable *L. monocytogenes* in food is the culture-based plate counting method. However, the cultivation process is time-consuming, involves cumbersome operations, demands strict environmental and operational conditions, and potentially leads to the oversight of viable but nonculturable (VBNC) cells [14–16]. Although the automated systems for colony inoculation and counting methods [17,18] can significantly enhance operational efficiency, the substantial time required for bacterial isolation and cultivation remains unresolved. This significant time lag severely restricts the timely response capabilities of food safety regulation and industrial quality control. To overcome the drawback of the delayed results from culture-based counting methods, several rapid detection techniques have been developed and applied. Rapid detection methods often utilize dyes to achieve quick differentiation and quantification of viable and dead bacteria. For instance, flow cytometry is frequently combined with dyes such as SYTO9/PI [19] and TO/PI [20]. Meanwhile, methods based on Calcein AM [21] or tetrazolium salt reduction and color development, such as MTT [22] and WST-8 [23], as well as ATP bioluminescence methods based on luciferin and luciferase [24], have been developed and typically the fluorescence intensity or absorbance are measured using a hand-held fluorometer or a microplate reader. These methods then establish a standard curve accordingly to quantify bacterial concentration. The rapid detection techniques are faster and more convenient than traditional culture methods and enable high-throughput data acquisition. However, for the specific detection of bacterial species, viable and dead stains cannot achieve the clonal complexes (CCs) typing of strains, which is crucial for tracing infection sources, identifying dominant pathogenic lineages, and studying the epidemiological characteristics of strains. Therefore, we urgently need a detection method that integrates speed, high accuracy, the ability to distinguish CCs typing, and precise quantification of viable bacteria.

Here we develop a microfluidic droplet digital PCR (ddPCR)-based method to rapidly and accurately quantify bacteria with viable/dead distinguishing and CC typing (Fig. 1.). To demonstrate CC typing, we detect the most frequently detected typing in clinical *L. monocytogenes* isolates in China— clonal complex 87 (CC87) [25–27]. By integrating propidium monoazide (PMA) pretreatment with droplet image processing and counting analysis, this method innovatively achieves rapid identification and quantification of viable CC87 *L. monocytogenes*. This method achieves a quantification limit of 3.3×10^2^ CFU/mL and completes the entire process within 3 hours, demonstrating high sensitivity and speed. When applied in samples with mixed strains and mixed species, this method successfully achieves strain-specific differentiation and precise quantification of viable bacteria. Overall, this technology effectively overcomes the aforementioned technical bottlenecks, achieves rapid and accurate detection of viable bacteria with CC typing, and provides a reliable tool for risk monitoring of highly pathogenic subtypes.

**Fig. 1.**
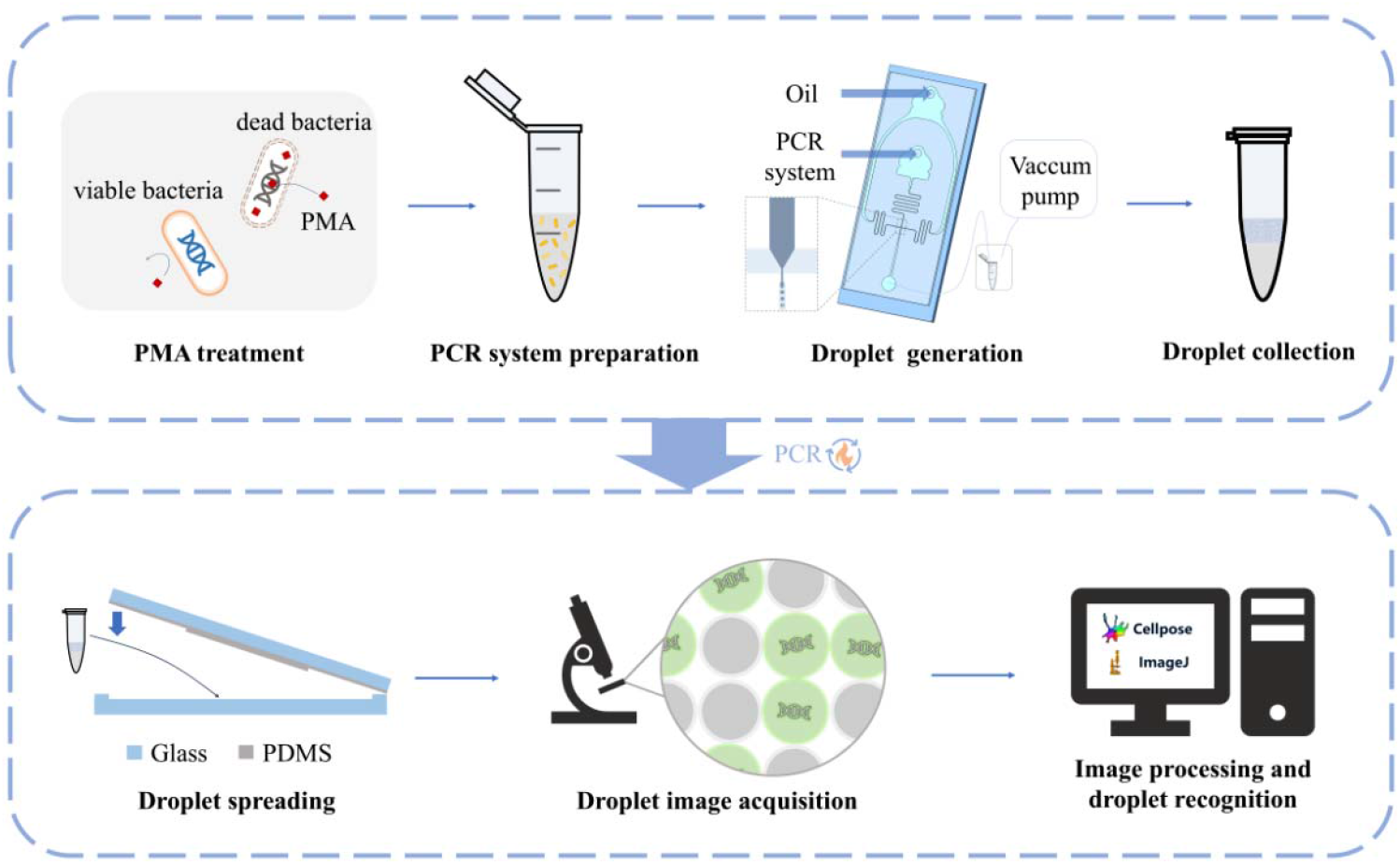
Procedure for rapid quantitative detection of viable *Listeria monocytogenes* clonal complex 87 based on microfluidic droplet digital PCR.

## 2. Materials and methods

### 2.1 Bacterial sample preparation and DNA extraction

Two strains of CC87 *L. monocytogenes*, MRL 300112 and MRL 300131, isolated in our laboratory, are selected as the test subjects. Frozen stocks of bacteria are maintained in Tryptone Soy Yeast Extract Broth (TSB-YE; Qingdao Haibo Biotechnology Co., Ltd., Qingdao, China) with 50% glycerol at −80°C. Working stocks are stored at 4°C on Tryptone Soy Agar with 0.6% Yeast Extract (TSA-YE; Qingdao Haibo Biotechnology Co., Ltd., Qingdao, China) and are renewed monthly. Prior to each experiment, a single colony is inoculated into 10 mL of TSB-YE and incubated in a shaker at 180 rpm and 37°C for 16 ∼ 18 h until reaching the stationary phase, resulting in an initial bacterial suspension with a concentration of approximately 10^9^ CFU/mL, which serves as the working solution. Preparation of dead bacterial suspension: Take the aforementioned working solution and treat it at 100°C for 15 min to obtain dead bacterial cells of *L. monocytogenes*. The viability of the cells can be confirmed by spreading them on TSA-YE and incubating at 37°C for 24 h; the absence of colony growth indicates dead bacterial cells. DNA extraction is performed using a DNA extraction kit (Tiangen Biotech, Beijing, China).

### 2.2 Target DNA fragment, primers and probe

Based on the SNP comparison analysis of 348 different CC-type *L. monocytogenes* strains, including 12 CC87 *L. monocytogenes* as shown in Fig. S1 Specific sequence fragments of CC87 *L. monocytogenes* are selected based on SNP comparison for the design of primers and probe. These primers and probe are subjected to homology comparison analysis on the BLAST platform of NCBI to verify their specificity. After confirmation, the synthesis is commissioned to Sangon Biotech (Shanghai, China) Co., Ltd.

### 2.3 Optimization of PCR reaction conditions

The amplification procedure and system are optimized through qPCR. The combination showing the lowest Ct value and the highest Rn value in qPCR results is considered optimal, which could enhance the amplification efficiency in subsequent ddPCR and increases the fluorescence intensity of positive droplets, thereby facilitating the distinction between positive and negative droplets. The qPCR reaction system was 20 μL, comprising qPCR premix (Sangon Biotech, Shanghai, China), 10 μM primers, 10 μM probe, ddH2O, and template DNA. The qPCR amplification procedure was as follows: 94°C for 10 min; 40 cycles of 94°C for 5s and 60°C for 30s, with fluorescence signal collection. Using CC87 *L. monocytogenes* (MRL 300112) as the positive template, the annealing temperature was optimized, and the usage amounts of primers and probe were screened. The procedure is tested at eight different annealing temperatures: 50°C, 52°C, 54°C, 56°C, 58°C, and 60°C, using the recommended primers/probe concentration of 200/100 nM. At the optimal annealing temperature, the final concentration ratios of primers/probe are set at 200/300 nM, 200/250 nM, 200/200 nM, 200/150 nM, 200/100 nM, and 200/50 nM.

### 2.4 Optimization of PMA treatment

Dissolve 1 mg of PMA (Aladdin, Shanghai, China) in 980 μL of distilled water to obtain a 2 mM stock solution, which is then set aside. Accurately measure a certain volume of the 2 mM PMA and add it to the sample to achieve a specific final concentration of PMA in the PMA-sample mixture. Thoroughly mix the sample and incubate it in a dark room for 15 minutes to allow PMA to penetrate dead cells and attach to DNA, ensuring full penetration of PMA into both dead and viable cells. After the dark treatment, place the sample in the PMA-Lite™ LED Photolysis Device (Biotium) and expose it to LED light (460 nm) for 15 minutes to induce the device of PMA with DNA from dead cells. After crosslinking, centrifuge the PMA-sample mixture at 10,000 rpm for 1 min, wash it twice with PBS (Adamas, Shanghai, China) to collect bacterial cells, and subsequently use it for DNA extraction and PCR. Dead and viable bacterial suspensions of *L. monocytogenes* are mixed with different volumes (0, 2.5, 5, 7.5, 10, 12.5, 15, 17.5, and 20 μL) of PMA solution to obtain PMA-sample mixtures with varying final PMA concentrations (0, 10, 20, 30, 40, 50, 60, 70, and 80 μM). These PMA-sample mixtures are pre-treated as described above, followed by bacterial DNA extraction using the aforementioned method, and finally subjected to quantitative fluorescent PCR analysis. Observe the test results to analyze and obtain the optimal PMA treatment conditions.

### 2.5 Design and fabrication of the microfluidic chip

Design the pattern of the droplet generation chip using AutoCAD 2019. Use SU-8 3025 as the photoresist and a silicon wafer as the substrate to fabricate the mold through MicroWriter ML3 maskless lithograph (Durham Magneto Optics, UK). Development is completed by immersion in 1-methoxy-2-propyl acetate (Sigma-Aldrich). After manufacturing the silicon wafer mold, the next step is to create the microfluidic chip using polydimethylsiloxane (PDMS, Sylgard184, Dow Coming, USA). Mix the PDMS base and curing agent in a 10:1 ratio, stir evenly, and then place it in a vacuum chamber to remove bubbles. Pour the mixture onto the silicon wafer wrapped in tin foil and place it on a hot plate at 90°C for 2 hours. Use tweezers to peel the obtained PDMS from the mold, cut it to the appropriate size, and use a puncher to create holes at the reserved locations on the chip. Place the cleaned glass slide, which has been washed with an ultrasonic cleaner, and the cut PDMS into a plasma cleaner for cleaning and bonding. After bonding, a complete microfluidic droplet generation chip is obtained.

### 2.6 ddPCR method

According to the optimized reaction protocol and primers and probe concentrations from qPCR, ddPCR experiments are conducted. The PCR reaction mixture serves as the aqueous phase, and fluorinated oil (ThunderBio Innovation Limited, Hangzhou, Zhejiang, China) is used as the oil phase. The experiment employs a droplet generation method based on negative pressure formed by a vacuum pump, with the PCR reaction system serving as the aqueous phase and the droplet generation oil as the oil phase. Driven by the negative pressure, the oil and aqueous phases enter the cross-shaped channel of the droplet generation chip, where the aqueous phase forms highly monodisperse droplets under the shear force of the oil phase. Droplets are generated and collected in PCR tubes, followed by amplification. After spreading the droplets using a homemade droplet spreading device, fluorescence images of the droplets are acquired using the Olympus IX83 inverted fluorescence microscope (Olympus, Japan). Droplet counting and size identification are performed using Cellpose [28] and ImageJ. Cellpose is utilized for droplet recognition and segmentation, followed by quantification using ImageJ. We fabricate the droplet spreading device through the following process: First, we tightly adhere a 50 μm-thick PDMS film (Shenzhen, China) to the center of a disposable petri dish. Subsequently, we mix epoxy resin AB glue at a 4:1 weight ratio, pour it into the petri dish, and let it cure overnight at room temperature on a level surface. After the mold fully cures, we peel it from the petri dish with the PDMS film-imprinted groove facing upward. Next, we slowly pour the prepared PDMS solution (following Method 2.5) onto the mold surface and cover it with a glass slide, taking care to avoid bubble formation during this step. We then place the entire assembly in an 80□□ oven for overnight curing. Finally, we separate the glass slide with the cured PDMS—which serves as the upper cover of the device —from the mold and assemble it with a glass counting chamber (Wuxi, China) that functions as the base component.

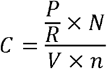

where *C* is the logarithm of the bacterial concentration (CFU/mL), *P* is the number of positive reaction units, *R* is the total number of reactions units, *N* is the total number of droplets generated by the system, *V* is the volume of bacterial solution added to the system (mL), *n* is concentration factor of the bacterial solution.

### 2.7 Anti-interference evaluation

In this study, two CC87 *L. monocytogenes* (MRL 300131 and MRL 300112) isolated from food and clinical samples in our laboratory are selected as target detection strains, while the reference strain *L. monocytogenes* EGD-e and the non-pathogenic *L. innocua* ATCC 33090 are used as interference strains to evaluate the anti-interference performance of the ddPCR method. All strains are cultured in TSA-YE to the stationary phase as described in Section 2.1, and the bacterial suspension concentration is adjusted to approximately 10□ CFU/mL. Subsequently, the CC87 target strains are mixed with each interference strain at ratios of 1:0, 1:1, and 0:1, respectively, to systematically evaluate the detection specificity and anti-interference capability of ddPCR under mixed bacterial conditions.

### 2.8 PMA-ddPCR method

After treatment under the PMA processing conditions optimized in Section 2.4, detection is performed using the ddPCR method described in Section 2.6. To verify the accuracy of the PMA-ddPCR method in detecting viable bacteria, a mixed bacterial suspension of dead and viable bacteria with a concentration of approximately 10□ CFU/mL is prepared according to the method described in Section 2.1. The mixed bacterial suspension is qualitatively verified by staining with the BacLight™ Live/Dead Bacterial Viability Kit (Servicebio, Wuhan, China) which contains two dyes, SYTO9 and propidium iodide (PI). Fluorescence images are acquired using the Olympus IX83 inverted fluorescence microscope (Olympus, Japan), and the images are processed using ImageJ software. Additionally, mixed samples of dead and viable bacteria with different proportions of viable bacteria (100%, 50%, 25%, and 0%) are prepared and subjected to comparative detection using both the plate counting method and the PMA-ddPCR method to further validate the quantitative performance of this method under conditions with varying proportions of viable bacteria.

### 2.9 Data acquisition and analysis

Data acquisition and analysis of fluorescence images are performed on MATLAB and OriginPro 2021. All experiments are repeated three times. Data are presented as mean ± standard deviation (SD). Significance analysis (ANOVA) is conducted using GraphPad Prism 8.0.2 software, with a significance level of p < 0.05. Data processing and analysis are carried out using OriginPro 2021.

## 3. Results

### 3.1 Rapid generation and identification of uniform droplets

We adopt a flow-focusing dropmaker design [29] to rapidly generate uniform droplets. In brief, the design consists of an oil phase inlet, an aqueous phase inlet, a droplet generation nozzle, and a droplet outlet. The complete design and image of the chip are shown in Fig. 2a. We employ the suction droplet technique, driving the liquid flow by applying a vacuum at the outlet, thereby enabling simple and rapid droplet generation. We achieve a generation rate of approximately 2000 droplets per second per device. Notably, this straightforward suction droplet scheme supports parallel droplet generation (Fig. 2b.), further enhancing the efficiency of droplet production. The generated droplets are uniform in size and can be quickly analyzed, with the entire detection process completed in approximately 3 hours. We employ the Cellpose image processing technique, which accurately identifies droplets in bright-field images (Fig. 3a-f.). We analyze the sizes of a total of 9260 droplets in three independent experiments. The average diameter of these droplets is 20.84 μm, with a coefficient of variation (CV) of 7.37% (Fig. 3g.), demonstrating a high degree of size uniformity.

**Fig. 2.**
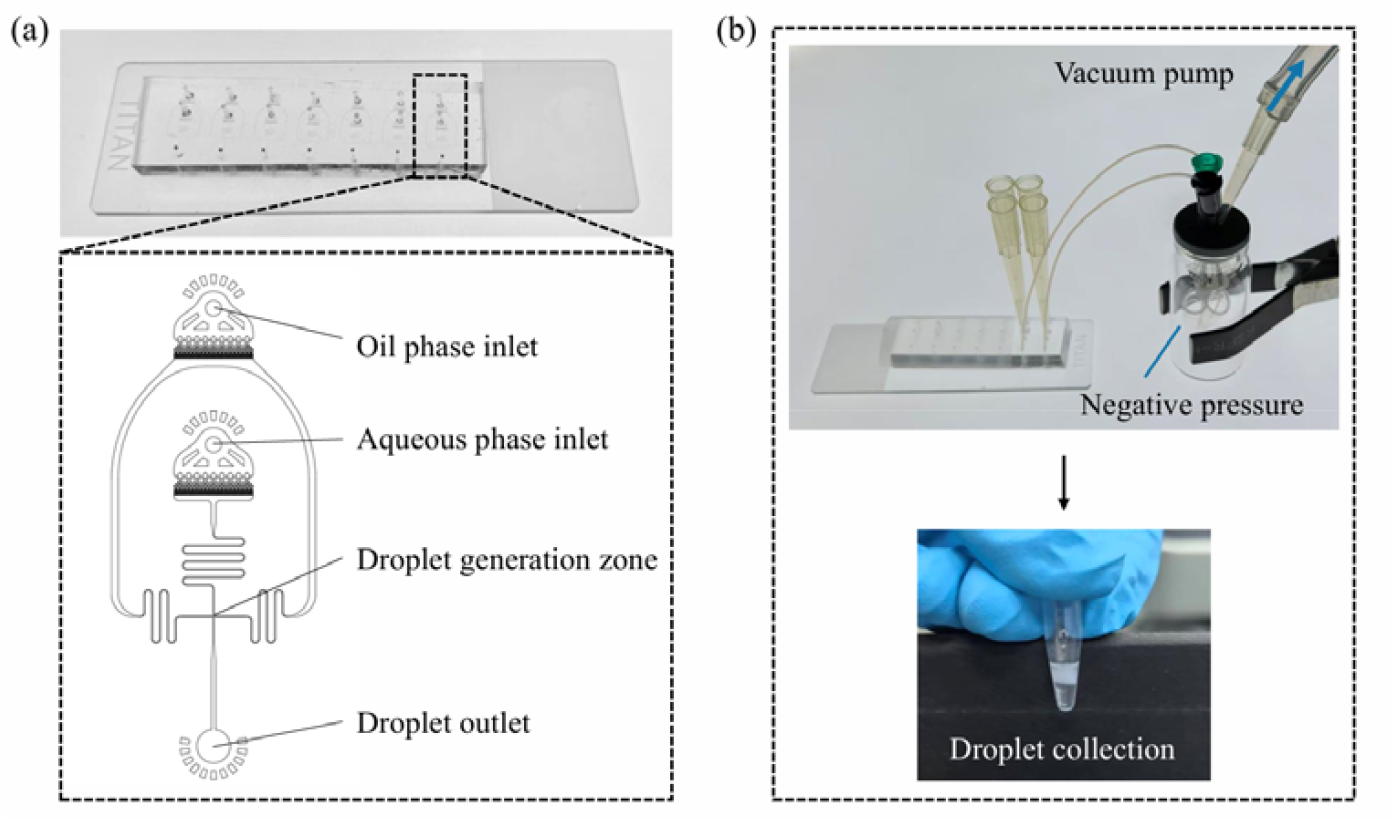
(a) Physical diagram and schematic diagram of the droplet generation chip structure; (b) Droplet generation device.

**Fig. 3.**
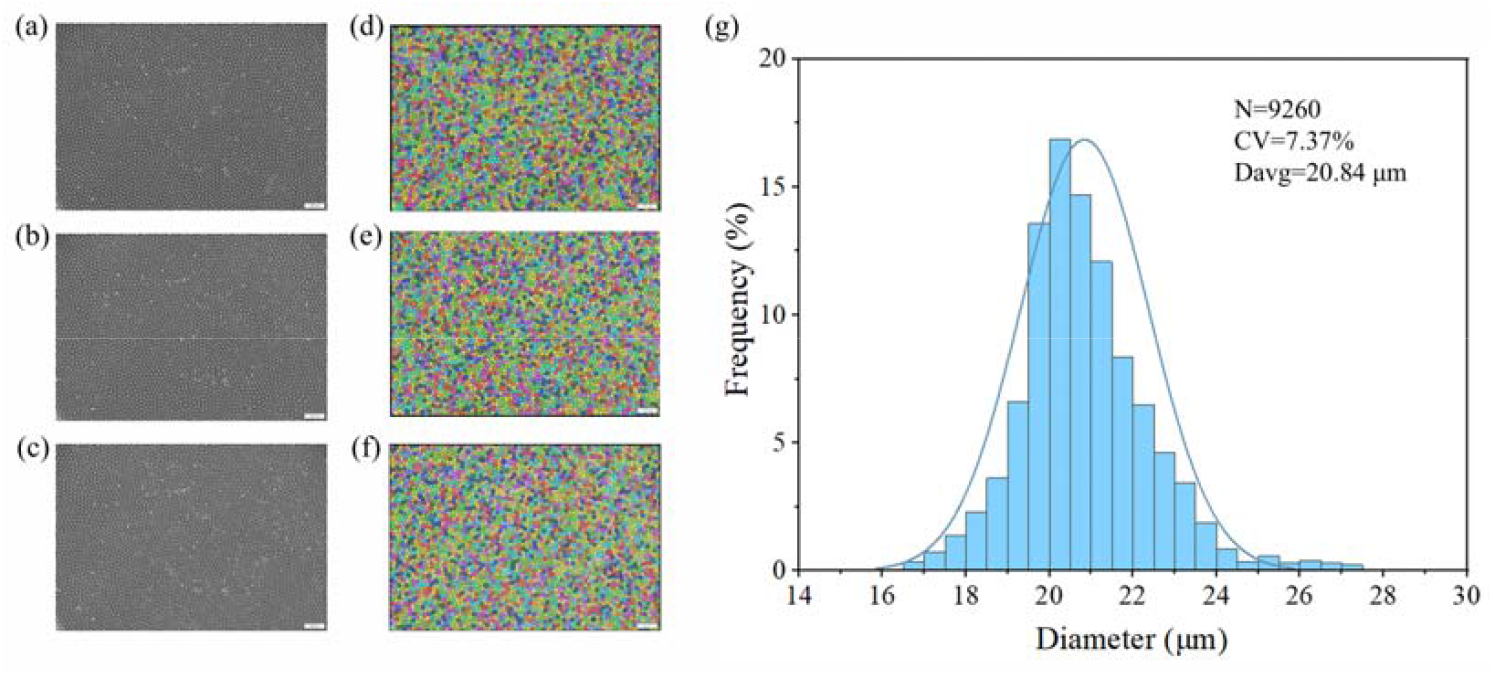
(a) - (c) Bright-field images of monodisperse droplets. (d) - (f) Fluorescence images of the same droplets after amplification, with false colors assigned by Cellpose image processing software to indicate successful identification. (g) Droplet diameter distribution, showing high uniformity. The average diameter is 20.84 μm with a coefficient of variation (CV) of 7.37%.

### 3.2 Developing a rapid ddPCR quantification method

We designed primers and probe (Table S1) for the successfully screened CC87-specific sequences, and their high specificity was validated through Blast and qPCR experiments (Fig. S2). The annealing temperature and primers/probe concentrations are optimized via qPCR, and the optimized parameters are applied to ddPCR quantitative analysis. The results show that the Ct value is the lowest and the fluorescence intensity Rn is the highest at an annealing temperature of 50°C (Fig. S3), thus 50°C is determined as the optimal annealing temperature. At a primers/probe concentration of 100/50 nM, the Ct value is the lowest, and there is no significant difference in Ct values among the three groups of 100/50, 200/50, and 150/50 nM (p > 0.05); while the Rn value reaches its highest at 300/50 nM, and there is no significant difference in Rn values among the four groups of 300/50, 200/50, 300/100, and 250/50 nM (Fig. S4). Considering both Ct values and Rn performance, the optimal primers/probe concentration is ultimately selected as 200/50 nM. This condition ensures amplification efficiency while significantly enhancing the differentiation between positive and negative droplets, and also takes into account the economy and stability of the reaction.

With optimized reaction mixture, the ddPCR system accurately quantify bacteria over a wide concentration range. We test the PCR system with a colony count range of approximately 10^2^ to 10^8^ CFU/mL as the aqueous phase. After droplet generation and amplification via a microfluidic chip, the proportion of positive droplets is calculated through fluorescence imaging analysis to determine the bacterial concentration. The ddPCR detection results exhibit a high degree of consistency with the traditional plate counting method (R^2^ = 0.9945, Fig. 4. and Table S2), indicating that this method possesses excellent linear quantitative capabilities, with a lower limit of quantification (LoQ) reaching 3.3 × 10^2^ CFU/mL.

**Fig. 4.**
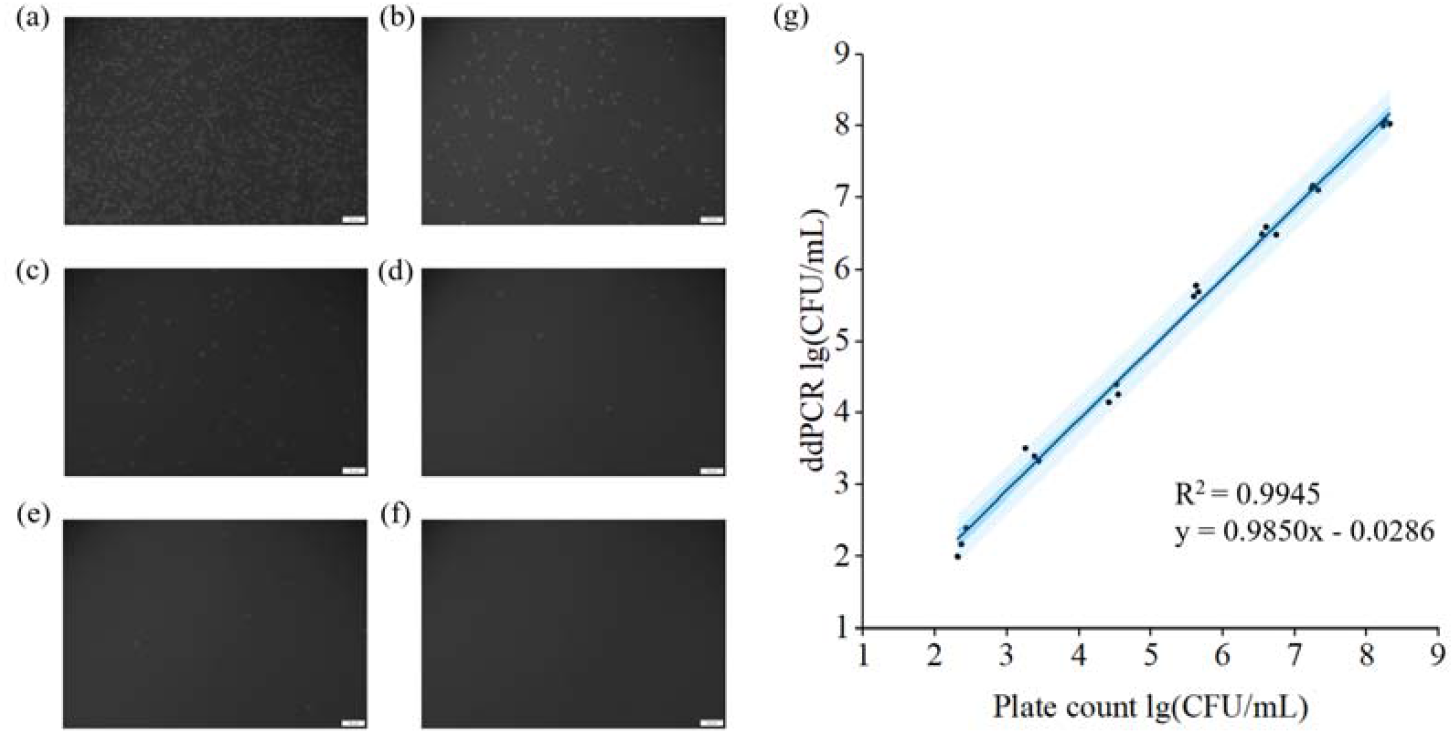
Quantitative detection of CC87 *L. monocytogenes* using ddPCR method. Successfully captured fluorescence field droplet images (a)-(f) under an inverted fluorescence microscope, with corresponding bacterial solution concentrations of (a) 10^8^ CFU/mL, (b) 10^7^ CFU/mL, (c) 10^6^ CFU/mL, (d) - (e) 10^5^ CFU/mL, and (f) negative control with ddH2O replacing the bacterial solution; (g) Correlation between bacterial concentrations determined by the ddPCR method and the plate counting method. The results show a high degree of consistency (R^2^ = 0.9945), indicating the accuracy of the ddPCR assay.

### 3.3 Specific Detection and Quantification of the CC87 *L. monocytogenes* by ddPCR

In the context of complex mixed microbial communities, the ddPCR method established in this study is capable of accurately identifying and quantifying CC types via primer specificity. By mixing the CC87 strains (MRL 300131, MRL 300112) with competitive non-target strains (*L. monocytogenes* EGD-e, *L. innocua* ATCC 33090) in different ratios, the anti-interference performance of ddPCR was evaluated. The results show that even under mixed bacterial conditions, the detection specificity of this method for the CC87 strains shows no significant difference compared to pure culture (p > 0.05), confirming its excellent specificity and anti-interference capability (Fig. 5.). This reproducible result confirms the anti-interference capability and specificity of the detection system, providing a reliable technical tool for addressing the monitoring challenges of specific subtypes in complex samples.

**Fig. 5.**
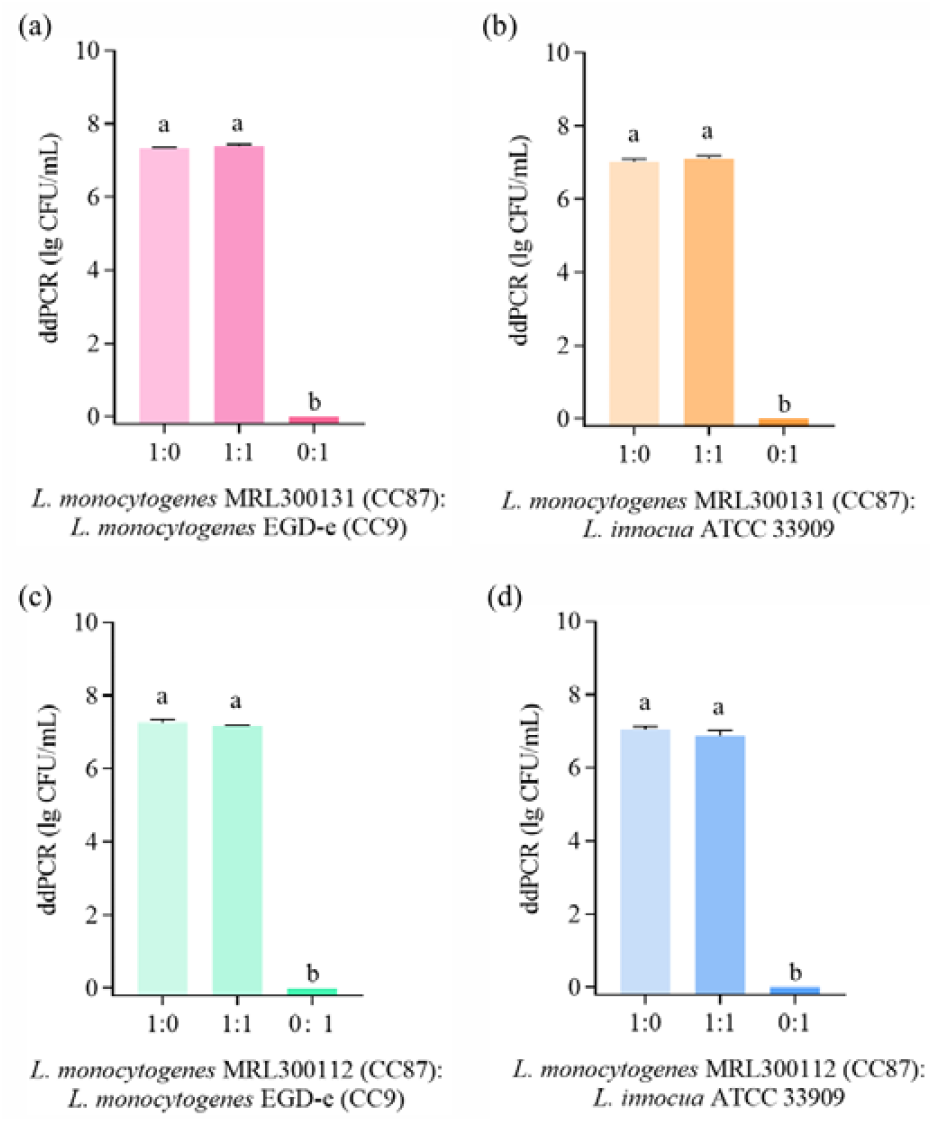
Precise quantification of CC87 *L. monocytogenes* by ddPCR. *L. monocytogenes* MRL300131 mixed with (a) *L. monocytogenes* EGD-e and (b) *L. innocua* ATCC 33909; *L. monocytogenes* MRL300112 mixed with (c) *L. monocytogenes* EGD-e and (d) *L. innocua* ATCC 33909. Same lowercase letters on the bars represent no significant differences, while different letters represent significant differences among treatments (p > 0.05 was non-significant).

### 3.4 Accurate differentiation and quantification of viable CC87 *L. monocytogenes* using the PMA-ddPCR method

Because dead bacteria cause false positive results in ddPCR, this study integrates PMA staining in the overall protocol to effectively remove dead bacteria signal and specifically quantifies viable bacteria. We perform systematic optimization of PMA treatment conditions and find that as the PMA concentration (20 ∼ 80 μM) increases, the Ct value of dead bacteria significantly increases, while the amplification of viable bacteria remains unaffected (p > 0.05; Fig. S5a), indicating that PMA selectively binds to dead bacterial DNA and inhibits its amplification, thus determining 20 μM as the optimal PMA treatment concentration. Optimization experiments for dark incubation time show that when the dark incubation time is ≥ 2 min, there is no significant difference in Ct values between viable and dead bacteria (p > 0.05; Fig. S5b), hence 2 min is selected as the optimal dark incubation time. The analysis of the impact of different illumination durations on the amplification efficiency of viable and dead bacteria reveals that within the range of 0 to 30 minutes, the amplification of viable bacteria is unaffected; in contrast, the Ct value of dead bacteria significantly increases as the illumination time increases from 0-15 minutes (Fig. S5c). Between 15 and 30 minutes, the Ct value of dead bacteria shows no significant change (p > 0.05), and the amplification of viable bacteria remains stable (p > 0.05). To maximize the elimination of interference from dead bacteria amplification and to minimize the processing time, 15 minutes is ultimately determined as the optimal illumination duration. The dCt values of viable bacteria are generally close to 0 ± 1, indicating that PMA does not affect the DNA amplification of viable bacteria; whereas the dCt values of dead bacteria are greater than 8, equivalent to a removal rate of approximately 99.6% of dead bacterial DNA. Additionally, we find that using different resuspension/washing solutions in PMA treatment affects its efficacy. Among the three resuspension/washing solutions, only PBS meets the requirements (Fig. S5d), maximizing the inhibition of dead bacterial amplification without affecting the amplification of viable bacteria. The BacLight™ Viability Kit is used for qualitative validation of the mixed bacterial suspension (Fig. 6a-c.). Fluorescence microscopy analysis shows green fluorescence for viable bacteria and red fluorescence for dead bacteria, providing a visual representation of the composition of viable and dead bacteria in the sample. It is noteworthy that in mixed bacterial suspension samples with different ratios, there is no significant difference in the quantitative detection results of viable bacteria between the PMA-ddPCR method and the plate counting method (p > 0.05; Fig. 6d. and Table S3). This result indicates that the PMA-ddPCR method can accurately distinguish and quantify viable bacteria, effectively excluding the interference of dead bacterial DNA, and achieving specific quantitative detection of viable CC87 *L. monocytogenes*.

**Fig. 6.**
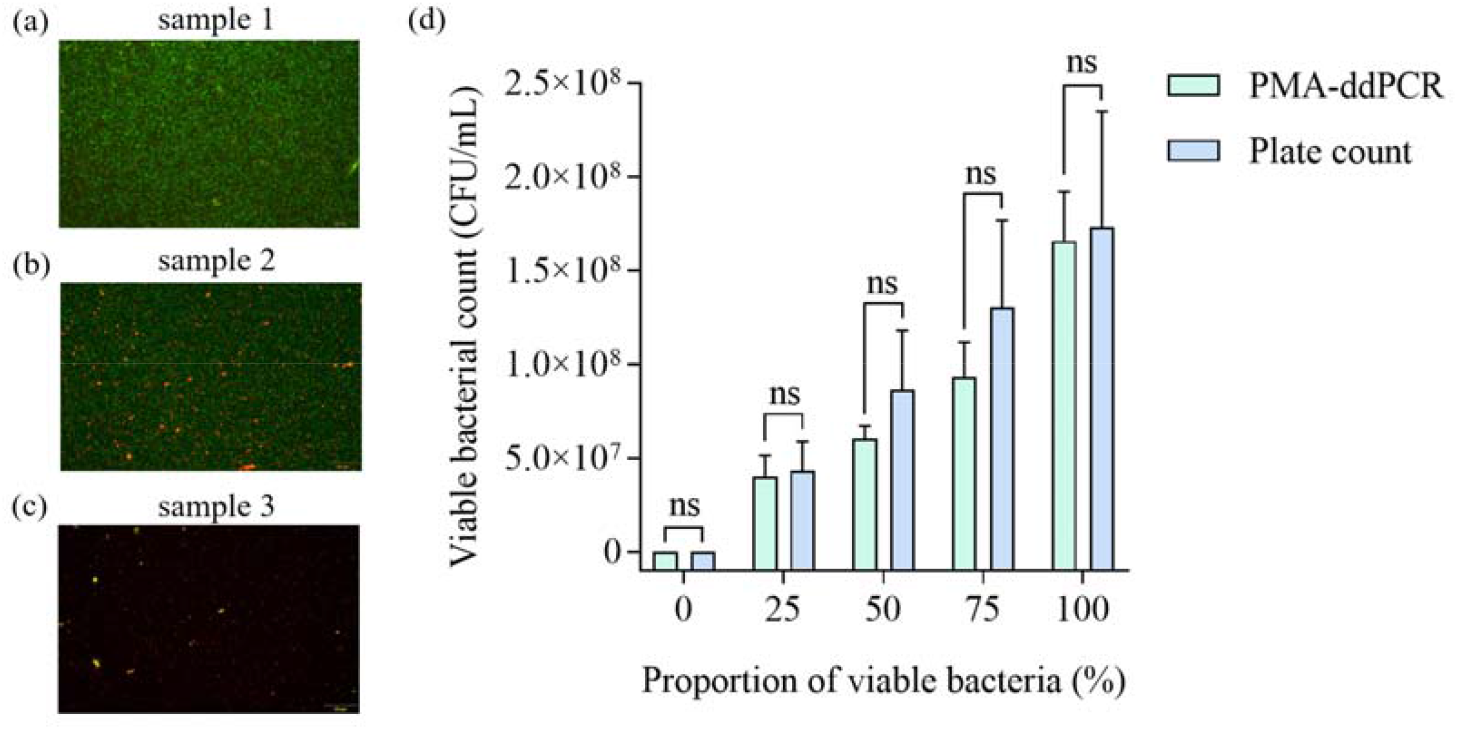
Evaluation and detection of CC87 *L. monocytogenes* bacterial suspension. (a)-(c) Fluorescence images of three distinct samples after staining with the SYTO9/PI Live-Dead viability kit. Viable bacteria fluoresce green, and dead bacteria fluoresce red. The samples are prepared as follows: Sample 1: A 1:10 dilution of a stationary-phase culture in PBS (∼10□ CFU/mL). Sample 3: Heat-inactivated Sample 1. Sample 2: A 1:1 (v/v) mixture of Sample 1 and Sample 3. (d) Quantification of viable bacteria in mixed suspensions with varying proportions of viable and dead cells, as determined by the PMA-ddPCR and plate counting methods.

## 4. Conclusion

In summary, we present a PCR method based on droplet microfluidics for sensitive and rapid detection of viable CC87 *L. monocytogenes* without nucleic acid extraction. The introduction of the PMA treatment procedure effectively eliminates the interference of dead bacteria. The droplet-based method has a LoQ for CC87 *L. monocytogenes* of 3.3 × 10^2^ CFU/mL within 3h. The method proposed in this study for the CC87 strains is an accurate and reliable quantitative technique for monitoring the CC87 strains under mixed bacterial conditions, and it can serve as a powerful tool for food safety assessment.

## Acknowledgments

This work was supported by Qinghai Provincial Department of Science and Technology Key Research and Development and Transformation Program (Grant No. 2025-QY-208), grants from the National Key Research and Development Program of China (No.2023YFA0915200, 2023YFA0915204) and the National Natural Science Foundation of China (32102095). This study was also supported by the equipment research and development projects of the Chinese Academy of Sciences (PTYQ2024YZ0010), the equipment research and development projects of the Chinese Academy of Sciences (PTYQ2024BJ0007), and the Science and Technology Commission of Shanghai Municipality Project (XTCX-KJ-2024-038).

## Supplementary information

Supplementary information associate with this article can be found.

