## Supplementary information for "Rapid and accurate quantification of viable *Listeria monocytogenes* with clonal specificity using microfluidic droplet digital PCR based technology"

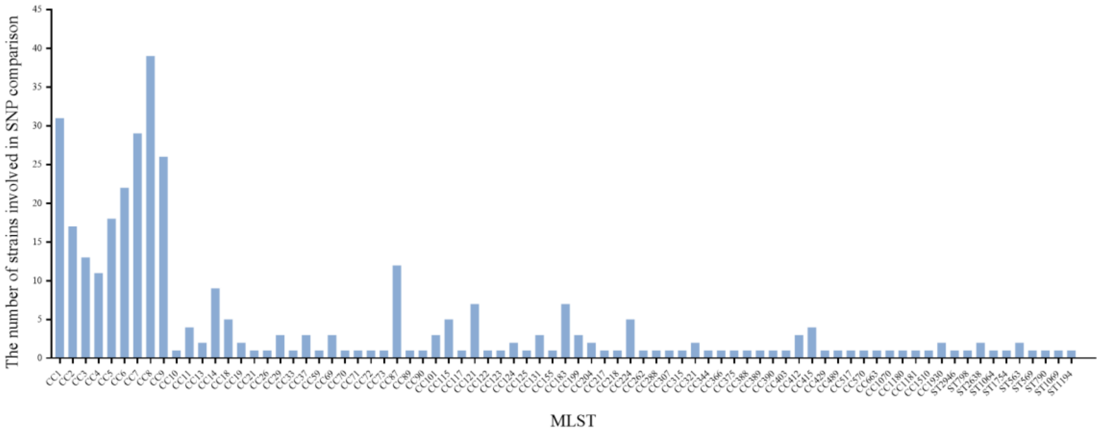


**Fig. S1.** MLST typing statistics of 348 *L. monocytogenes* involved in SNP comparison (including 12 strains of CC87 *L. monocytogenes*).

(a)

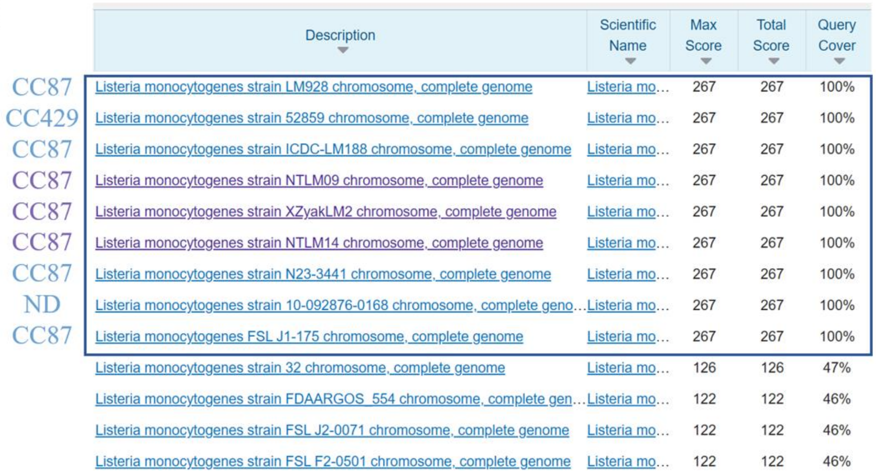

| | Description | Scientific Name | Max Score | Total Score | Query Cover |
|---|---|---|---|---|---|
| CC87 | Listeria monocytogenes strain LM928 chromosome, complete genome | Listeria mo... | 267 | 267 | 100% |
| CC429 | Listeria monocytogenes strain 52859 chromosome, complete genome | Listeria mo... | 267 | 267 | 100% |
| CC87 | Listeria monocytogenes strain ICDC-LM188 chromosome, complete genome | Listeria mo... | 267 | 267 | 100% |
| CC87 | Listeria monocytogenes strain NTLM09 chromosome, complete genome | Listeria mo... | 267 | 267 | 100% |
| CC87 | Listeria monocytogenes strain XZyakLM2 chromosome, complete genome | Listeria mo... | 267 | 267 | 100% |
| CC87 | Listeria monocytogenes strain NTLM14 chromosome, complete genome | Listeria mo... | 267 | 267 | 100% |
| CC87 | Listeria monocytogenes strain N23-3441 chromosome, complete genome | Listeria mo... | 267 | 267 | 100% |
| ND | Listeria monocytogenes strain 10-092876-0168 chromosome, complete geno... | Listeria mo... | 267 | 267 | 100% |
| CC87 | Listeria monocytogenes FSL J1-175 chromosome, complete genome | Listeria mo... | 267 | 267 | 100% |
| | Listeria monocytogenes strain 32 chromosome, complete genome | Listeria mo... | 126 | 126 | 47% |
| | Listeria monocytogenes strain FDAARGOS_554 chromosome, complete gen... | Listeria mo... | 122 | 122 | 46% |
| | Listeria monocytogenes strain FSL J2-0071 chromosome, complete genome | Listeria mo... | 122 | 122 | 46% |
| | Listeria monocytogenes strain FSL F2-0501 chromosome, complete genome | Listeria mo... | 122 | 122 | 46% |

(b)

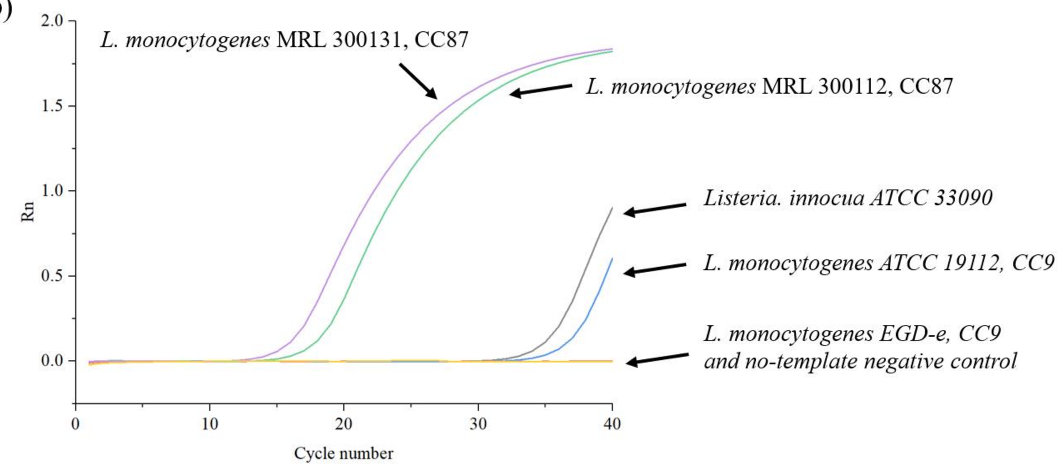


**Fig. S2.** Specificity validation of primers and probe. (a) BLAST alignment results of primers and probe on NCBI (https://www.ncbi.nlm.nih.gov/); (b) Specific test of qPCR for CC87 *L. monocytogenes* strains (non-target bacterial Ct values all >35, considered negative).

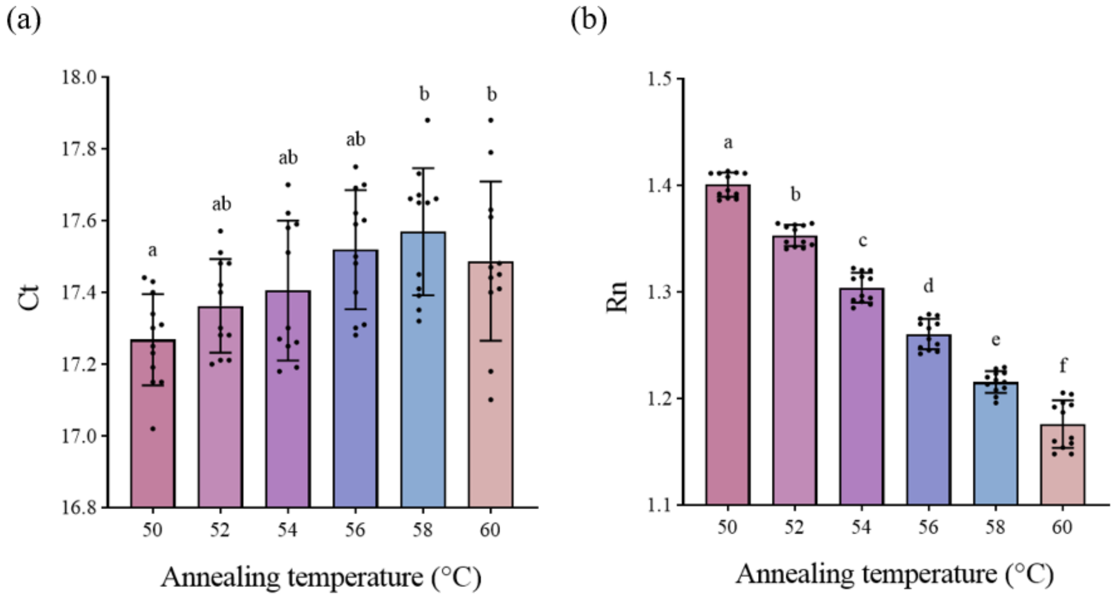


**Fig. S3.** Optimization of annealing temperature (a) Effect of Annealing Temperature on Ct Value; (b) Effect of Annealing Temperature on Rn Value (No significant difference in Ct between 50°C and 56°C, $p > 0.05$; Significant differences in Rn between 50°C and 60°C, $p < 0.05$).

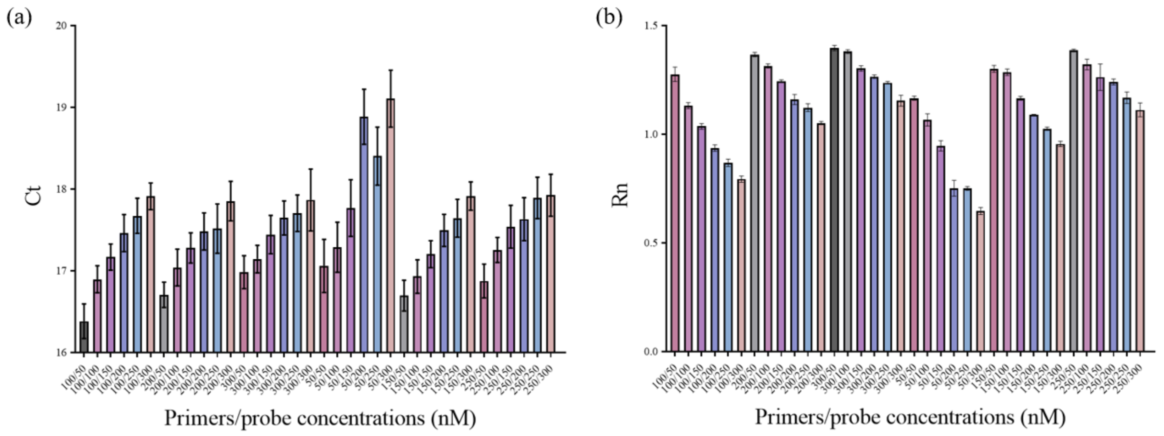


**Fig. S4.** Optimization of primers/probe concentration (a) Effect of primers/probe concentration on Ct values (the average Ct value is minimized at the primers/probe concentration marked in dark gray, with no significant difference compared to Ct values at concentrations marked in light gray, $p > 0.05$); (b) Effect of primers/probe concentration on Rn values (the average Rn value is maximized at the primers/probe concentration marked in dark gray, with no significant difference compared to Rn values at concentrations marked in light gray, $p > 0.05$).

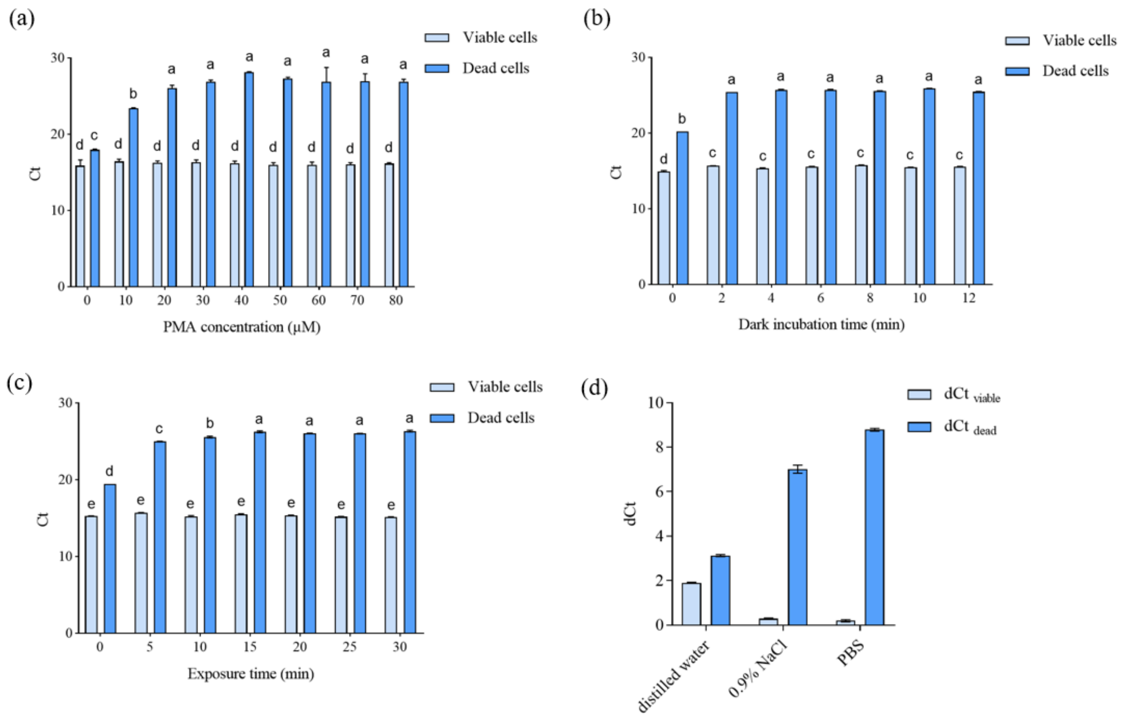


**Fig. S5.** Optimization of PMA treatment parameters (a) PMA concentration; (b) Dark incubation time; (c) Light exposure time optimization; (d) Effect of different resuspension/washing solutions on viable and dead bacteria during PMA treatment. dCt = Ct (PMA treated) - Ct (PMA untreated).

**Table S1** CC87 *L. monocytogenes* primers and probe

| **Type** | **Sequence (5'→3')** | **Amplicon length (bp)** |
|---|---|---|
| Forward primer | GCAAGGGCTCCAAGTAAGTAG | |
| Reverse primer | TGATGAGAACCCGCGTTAT | 144 |
| Probe | TCCCGGTTCCAAAGCTACCATGA | |

**Table S2** Quantification of CC87 *L. monocytogenes* strains bacterial loads using ddPCR across multiple orders of magnitude

| ddPCR lg (CFU/mL) | | | | | Plate count lg (CFU/mL) | Absolute error lg (CFU/mL) |
|---|---|---|---|---|---|---|
| **Parallel 1** | **Parallel 2** | **Parallel 3** | **Mean** | **CV/%** | | |
| 8.20 | 8.26 | 8.22 | 8.23 | 0.28% | 8.27 | -0.54 |
| 7.40 | 7.45 | 7.39 | 7.41 | 0.31% | 7.27 | 1.95 |
| 6.78 | 6.88 | 6.78 | 6.81 | 0.70% | 6.64 | 2.64 |
| 5.98 | 5.93 | 5.87 | 5.93 | 0.75% | 5.64 | 5.00 |
| 4.70 | 4.45 | 4.56 | 4.57 | 2.23% | 4.50 | 1.56 |
| 3.30 | 3.38 | 3.44 | 3.37 | 1.64% | 3.33 | 1.26 |
| 2.30 | 2.70 | 2.48 | 2.49 | 6.53% | 2.37 | 4.99 |

**Table S3** Quantification of CC87 *L. monocytogenes* strains bacterial loads using PMA-ddPCR across multiple orders of magnitude

| Proportion of viable bacteria/% | PMA-ddPCR results lg(CFU/mL） | | | | | Plate count results lg (CFU/mL) | PMA-ddPCR and plate count mean lg relative deviation/% |
|---|---|---|---|---|---|---|---|
| | Parallel 1 | Parallel 2 | Parallel 3 | Mean | CV/% | | |
| 0% | 0.00 | 0.00 | 0.00 | 0.00 | - | 0 | - |
| 25% | 7.64 | 7.44 | 7.69 | 7.59 | 1.46% | 7.64 | 0.61 |
| 50% | 7.75 | 7.84 | 7.75 | 7.78 | 0.52% | 7.94 | 2.03 |
| 75% | 7.87 | 8.04 | 7.98 | 7.96 | 0.92% | 8.11 | 1.87 |
| 100% | 8.22 | 8.14 | 8.28 | 8.22 | 0.69% | 8.24 | 0.28 |
